# Targeting the PI3K/AKT pathway overcomes enzalutamide resistance by inhibiting induction of the glucocorticoid receptor

**DOI:** 10.1101/783803

**Authors:** Remi M. Adelaiye-Ogala, Berkley Gryder, Yen Thi Minh Nguyen, Aian Neil Alilin, Adlai Grayson, Keith H. Jansson, Michael L. Beshiri, Supreet Agarwal, Jose Antonio Rodriguez-Nieves, Brian Capaldo, Kathleen Kelly, David J. VanderWeele

## Abstract

The PI3K-AKT pathway has pleiotropic effects, and its inhibition has long been of interest in the management of prostate cancer, where a compensatory increase in PI3K signaling has been reported following Androgen Receptor (AR) blockade. Prostate cancer cells can also bypass AR blockade through induction of other hormone receptors, in particular the glucocorticoid receptor (GR). Here we demonstrate that AKT inhibition significantly decreases cell proliferation through both cytostatic and cytotoxic effects. The cytotoxic effect is enhanced by AR inhibition and is most pronounced in models that induce compensatory GR expression. AKT inhibition increases canonical AR activity and remodels the chromatin landscape, decreasing enhancer interaction at the GR gene (*NR3C1*) locus. Importantly, it blocks induction of GR expression and activity following AR blockade. This is confirmed in multiple *in vivo* models, where AKT inhibition of established xenografts leads to increased canonical AR activity, decreased GR expression, and marked anti-tumor activity. Overall, our results demonstrate that inhibition of the PI3K/AKT pathway can block GR activity and overcome GR-mediated resistance to AR-targeted therapy. Ipatasertib is currently in clinical development, and GR induction may be a biomarker to identify responsive patients or a responsive disease state.

**SIGNIFICANCE:** Induced GR expression is compensatory for AR blockade and confers resistance to AR-targeted therapy. Here we show that inhibition of the PI3K/AKT pathway remodels the chromatin landscape, blocks the induction of GR expression and overrides enzalutamide resistance.

## INTRODUCTION

Suppression of androgen synthesis and androgen receptor (AR) activity via chemical castration and/or an AR antagonist is the mainstay of systemic prostate cancer therapy ^1^. While almost all patients have an initial favorable response, inevitably resistance develops and patients relapse with progressive disease ^2,3^. Multiple mechanisms can promote resistance, including reactivation of AR through amplification of the AR gene and/or an enhancer, mutation, or expression of AR variants ^4–7^. Expression of alternate hormone receptors with overlapping downstream targets can also promote resistance. Recent preclinical and clinical studies suggest the glucocorticoid receptor (GR) is the primary hormone receptor whose activity confers resistance to AR-targeted therapy ^8,9^. Resistance to AR-targeted therapy through induction of GR expression is the basis for multiple clinical trials combining inhibition of GR with enzalutamide (NCT03674814, NCT02012296, NCT03437941).

There is also cross-talk between androgen signaling and PI3K/AKT pathway activation. The PI3K/AKT pathway is a complex, branching and looping signaling pathway that is involved in survival, proliferation, metabolism, and growth pathways ^10^. In prostate cancer, castration or AR antagonism has been shown to increase phosphorylation of AKT and downstream targets, and loss of PTEN, a negative regulator of the PI3K/AKT pathway, correlates with decreased AR activity ^11^. Ipatasertib is a potent, pan-AKT inhibitor that binds AKT in an ATP-competitive manner, consequently disrupting its effect on downstream targets ^12,13^. It has activity in combination with the CYP17A1 inhibitor abiraterone plus prednisone in metastatic castrate-resistant prostate cancer (mCRPC)^14^. Given encouraging preliminary data, there are several ongoing clinical trials assessing the therapeutic benefit of ipatasertib in combination with other antineoplastic agents, including in prostate cancer ^15–17^; however, the optimal setting for AKT inhibition is unknown.

Here we demonstrate that inhibition of AKT has a cytostatic effect on prostate cancer cells despite increasing AR activity and is cytotoxic in combination with AR antagonism. Moreover, among the effects of PI3K/AKT pathway inhibitors, they block the induction of GR seen in response to AR blockade, which is associated with remodeling of the chromatin landscape and effects on transcriptional regulation. In *in vivo* models of established tumors, AKT inhibition enhances canonical AR activity, blocks GR expression, and demonstrates marked antitumor activity.

## MATERIALS AND METHODS

### Cell Lines and Reagents

LNCaP and C4-2 were purchased from ATCC (ATCC.com). LAPC4 and LREX were provided by Dr. Sawyers, Memorial Sloan Kettering Cancer Center. LNCaP, C4-2, 22Rv1 and LREX cells were cultured in RPMI1640+GlutaMAX (Gibco, Life Technologies), 10% FBS (Gibco, Life Technologies), 1% Penicillin/Streptomycin (Corning). For androgen deprivation *in vitro* studies, 10% charcoal stripped serum (CSS) (Gibco, Life Technologies) was used in place of 10% FBS. All cell lines were tested for mycoplasma contamination (PCR Mycoplasma Detection; primer sequence in **Supplementary Table 1**) every 6 months. LuCaP PDX models (provided by Dr. E. Corey and Dr. R. Vessella, Washington University) were grown in organoid culture in advanced DMEM/F12 media with supplements, as previously published ^18^. For *in vitro* studies, ipatasertib (Chemietek) and enzalutamide (Selleckchem) were dissolved in DMSO. CellTitre Glo Assay (Promega) was used to assess cell viability. For *in vivo* studies, ipatasertib (Division of Cancer Treatment and Diagnosis, NCI Developmental Therapeutics Program) and Enzalutamide (DCTD, NCI DTP) were dissolved in 1:1 of labrasol (Gattefosse) to PEG400 (Sigma).

### Western Blot Analysis

Cell lines and xenograft tumors collected at the end of treatment (EOT) time point were lysed using standard protocol. Lysates generated were used to perform immunoblot as previously described ^19^. Primary antibodies are diluted 1:1000 and from Santa Cruz (AR) or Cell Signaling Technology (AKT, p-AKT(Thr308), p-AKT (Ser483), GR, P70S6K, p-P70S6K, 4EBP1, p-4EBP1, and GAPDH). Secondary antibodies against rabbit (1:5000, EMD Millipore) or mouse (1:5000; EMD Millipore) were used followed by band detection using Amersham ECL Prime western blotting detection reagent (GE Lifesciences) per manufacturer’s instructions and visualized using ChemiDoc Touch Imaging system (Bio-Rad). Quantitative measurements of western blot analysis were performed with ImageJ and GraphPad Prism8 software.

### Quantitative RT-PCR (qRT-PCR) Analysis

mRNA extracts from cell lines and xenograft collected post EOT time points were used to perform qRT-PCR as previously described ^19^ for detection of KLK3, NKX3-1, and NR3C1 (**Supplemental Table 1**). PCR was performed on a StepOnePlus Real-Time PCR System (Applied Biosystems). PCR arrays were used to evaluate expression of a panel of 84 genes putatively regulated by GR (PAHS-154ZA) or by AR (PAHS-142ZA) (Qiagen). For the AR array, seven genes were removed from further analysis because their expression was increased by enzalutamide.

### RNA-Seq Analysis

RNA was extracted, quantified and profiled for quality using a Bioanalyzer (Agilent). Poly-A enriched and Illumina barcoded libraries were prepared and sequenced on a NextSeq 500 (Illumina), to a depth of 30 million base pairs (150 bp paired end) at the Illumina Sequencing Facility, Center for Cancer Research, NIH Frederick Campus. Paired-End RNA-seq reads were mapped to UCSC reference genome for hg19 with STAR (https://github.com/alexdobin/STAR), then Transcripts per Million (TPM) was calculated for each gene using RSEM (https://deweylab.github.io/RSEM). GSEA was performed using pre-ranked gene lists of log2 fold change comparison of TPM values between each treatment and the DMSO control. GSEA data visualization was performed using custom R scripts (available here: https://github.com/GryderArt/VisualizeRNAseq/).

### ChIP-Seq Analysis

Chromatin was isolated from cells using ChIP-IT High Sensitivity Kit (Active Motif) according to the manufacturer’s protocol. Sheared chromatin was incubated with antibody against H3K27ac (Active Motif). Protein-DNA bound complex was immunoprecipitated, followed by reverse-cross linking and DNA purification. ChIP-seq samples were pooled and sequenced on NextSeq (Illumina) using Accel-NGS 2S Plus DNA Library Kit (low input) (Swift) and single-end sequencing. All the samples have percent of Q30 bases above 92%. All the samples have yields between 15 and 57 million pass filter reads. Samples were trimmed for adapters using trimmomatic software before the alignment. Single-End reads were mapped to hg19 with BWA, TDFs were made for visualization in IGV using igvtools count (https://software.broadinstitute.org/software/igv/igvtools_commandline). MACS2 was used to perform peak calling with a threshold of 1E-7. HOMER was used to identify motif enrichment (http://homer.ucsd.edu) at sites of H3K27ac, and p-values were visualized in GraphPad Prism. Chromatin folding was inferred from analysis of public HiC data ^20^ (https://aidenlab.org/juicebox/).

### Gene Overexpression and Knockdown Assays

For GR (*NR3C1*) overexpression transfections, cells were transfected with 1-2μg NR3C1(Myc-DDK-tagged) expressing cDNA (OriGene Technologies, Inc.) using TurboFectin 8.0 transfection reagent according to manufacturer’s protocol (OriGene Technologies, Inc.) in LREX and LNCaP cells. For gene silencing of AR and NR3C1 gene expressions LREX cells were transduced with lentivirus particles of 4 different sequences per target gene or a non-silencing sequence as control (OriGene Technologies, Inc.) according to the manufacturer’s guidelines.

### *In Vitro* Assays

#### 2D in vitro experiments

LNCaP and LAPC4 cells were plated in 24-well plates. Following overnight incubation cells received fresh media containing 10% charcoal stripped serum (CSS) and 1% penicillin/streptomycin. Cells were treated with either 10nM R1881 (Sigma), 100nM ipatasertib or combination of R1881 and ipatasertib. LREX, 22Rv1, and C4-2 cells were prepared as previously stated and treated with either 100nM ipatasertib, 2μM enzalutamide or combination of both ipatasertib and enzalutamide. Viable cells were determined using CellTitre Glo assay 24hrs to 168hrs post treatment in accordance with manufacturer’s instructions and absorbance read using NanoQuant Infinite M200 Pro reader (TECAN). In parallel, cells were plated in 10cm dishes and collected 48hrs after treatment for RNA and protein analysis.

### *In Vivo* Studies

#### Xenograft models

LuCaP 136 and LuCaP 147 are well characterized prostate cancer(PCa) PDX models ^21^ established at the University of Washington. LuCaP 136CR-N is a castrate resistant PCa developed from the parental LuCaP 136 model that has undergone several passages in castrate mice to select for castrate resistant tumors. All LuCaPs were validated using short tandem repeat (STR) analysis.

#### Tumor implantation

All preclinical *in vivo* experiments were performed in accordance with an NCI Animal Care and Use Committee approved protocol. Six-week-old Athymic Nude-*Foxn1*^*nu*^ male mice were housed in a sterile, pathogen-free facility and maintained in a temperature-controlled room under a 12-hour light/dark schedule with water and food ad libitum. All mice were operated under sedation with oxygen and isoflurane. Ibuprofen and/or buprenorphine was administered post-surgery. Androgen sensitive prostate cancer xenograft (PDX) models LuCaP 136 or 147 were implanted subcutaneously under the left flank of intact mice. When tumors were established and reached an average volume of ∼ 500mm^3^, mice were randomized and placed in either control group or treatment groups (n=8/group for LuCaP 136, n=6/group for LuCaP 147). Mice received ipatasertib treatment (100mg/kg 5 days on 2 days off; oral gavage), androgen deprivation therapy (surgical castration) or a combination of both. For CRPC models LuCaP 136CR-N or LREX cells were implanted subcutaneously under the left flank of castrated mice. When tumors were established and reached ∼ 500mm^3^, mice were randomized and placed in either control group or treatment groups (n=8/group). Mice received ipatasertib treatment (100mg/kg 5days on 2 days off; oral gavage), enzalutamide (10mg/kg; 5days on 2 days off; oral gavage) or a combination of both. Tumor burden was assessed twice per week by caliper measurement of two diameters of the tumor (L × W = mm^2^) and reported as tumor volume ((L × W^2^)/2 = mm^3^). Body weights were assessed using a weighing scale and recorded in grams. Tumor tissues were excised, weighed and snap frozen for further analysis.

### Statistics

Statistical analysis was performed using GraphPad prism (version 7.0a). In brief, data analyses are expressed as the mean + standard error of mean (SEM) unless otherwise stated. Statistical significance where appropriate was evaluated using a two-tailed student t test when comparing two groups, or by one-way analysis of variance (ANOVA), using the student-Newman Keuls post-test for multiple comparison. A p value *p <0.05, **p < 0.01, ***p <0.001, was considered significant; ns= not significant, or otherwise stated.

## RESULTS

### The pan-AKT inhibitor ipatasertib decreases cell viability in an enzalutamide-resistant prostate cancer cell line

The PI3K/AKT signaling pathway is hyperactive in 50-60% of advanced prostate cancer ^22^, and the AKT inhibitor ipatasertib is currently in clinical development. We sought to determine the effects of ipatasertib (100nM) or enzalutamide in models representing castrate resistant prostate cancer (CRPC). We utilized 22RV1, which is known to harbor full length AR and ARV7, C4-2 (LNCAP/AR derived CRPC) and LREX (LNCAP/AR Resistant to Enzalutamide). Cells were exposed to enzalutamide (2 μM) or ipatasertib (100nM) in the setting of androgen deprivation. Response to enzalutamide was modest in 22Rv1 and C4-2 cell lines, as was response to ipatasertib (Fig. 1a, **1^st^ and 2^nd^ panel**). In contrast, LREX, which is poised to upregulate GR expression to achieve enzalutamide resistance^8^, was confirmed to be completely resistant to enzalutamide but had a marked response to ipatasertib (Fig. 1a, ***3^rd^ panel***). Consistent with its mechanism of action, the effect of ipatasertib on cell viability correlated with inhibition of AKT signaling, which is manifest as an increase in AKT phosphorylation (due to its ability to protect against phosphatases) and decrease in phosphorylation of downstream targets (Fig. 1b). To determine if ipatasertib would be of benefit in hormone sensitive prostate cancer (HSPC), we tested its effect alone vs. androgen withdrawal in the well-characterized HSPC cell lines LNCaP and LAPC4. Both models of HSPC were more sensitive to ipatasertib than to androgen withdrawal (**Extended Data Fig. 1a**). Western blot analysis confirmed ipatasertib inhibited AKT signaling.(**Extended Data Fig. 1b**). Taken together, these results indicate heterogenous response to ipatasertib as monotherapy. Amongst all 3 CRPC models tested, ipatasertib was most active in the enzalutamide resistant CRPC (LREX) cells.

**Fig. 1.**
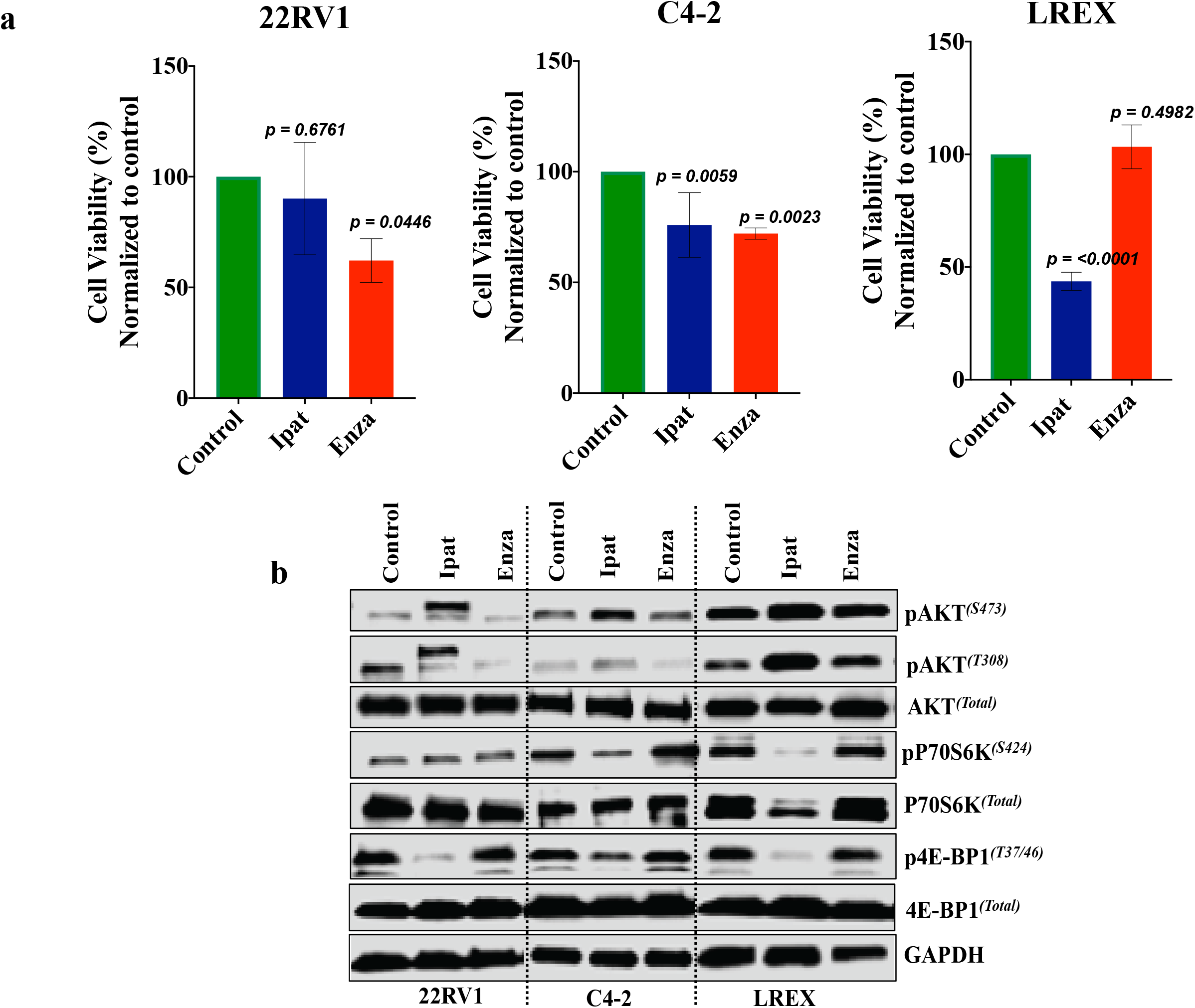
Ipatasertib as single agent is highly potent in enzalutamide resistant CRPC. **a**, Assessment of ipatasertib as single agent across PCa modeling castration resistance (22RV1 and C4-2), show marginal response to the pan-AKT inhibitor ipatasertib. LREX, which induces GR to achieve enzalutamide resistance, shows robust response to ipatasertib. **b**, Western blot indicates inhibitory effects on AKT1 phosphorylation and downstream targets. *Graphs are presented as mean +/− SD*.

### Ipatasertib induces cell cycle arrest as monotherapy and apoptosis in combination with enzalutamide

The LREX cell line was derived from LNCaP/AR with *in vivo* selection for enzalutamide resistance ^8,23^. We evaluated response to enzalutamide and ipatasertib alone and in combination in a dose dependent manner for 48 hours. We confirmed that LREX cells were resistant to enzalutamide at 10uM, with no difference in cell viability compared to DMSO treated cells (Fig. 2a). However, we observed response to ipatasertib single agent in a dose dependent manner which was enhanced when combined with single dose enzalutamide (Fig. 2a). Similarly, in a time dependent manner (24hrs to 168hrs of treatment) using single dose ipatasertib (100nM) and Enzalutamide, we again observed no response to enzalutamide (Fig. 2b). LREX cells continue to be sensitive to ipatasertib over time, with approximately 50% growth suppression. Combination of ipatasertib and enzalutamide led to almost complete growth suppression (Fig. 2b).

**Fig. 2.**
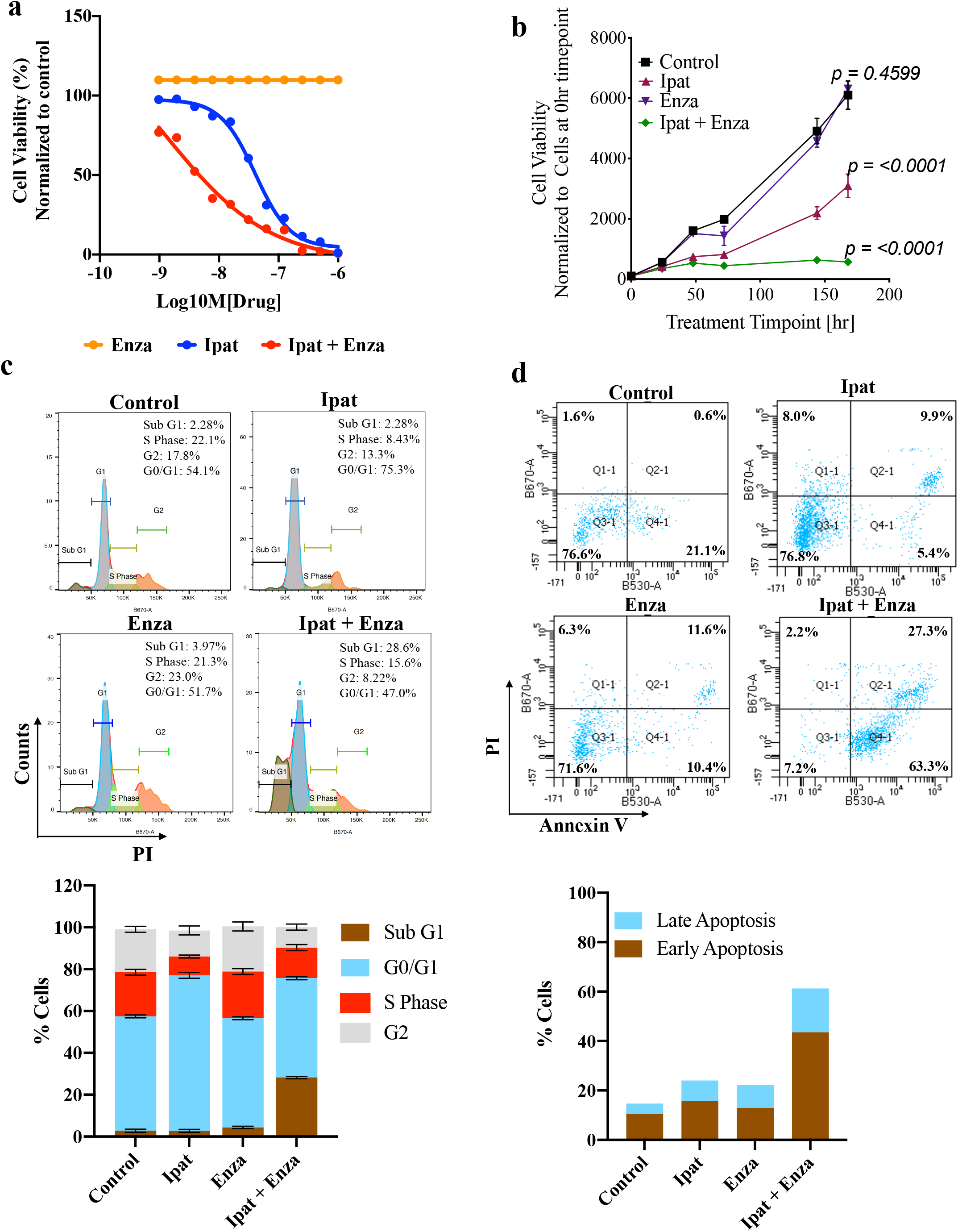
Ipatasertib overrides enzalutamide resistance, inducing cell cycle G0/G1 arrest and apoptosis. **a**, Cell proliferation assay with increasing concentrations of ipatasertib, single dose enzalutamide (2µM) or combination of both drugs shows the shift in cells treated with both ipatasertib and enzalutamide compared to ipatasertib alone. **b**, Time-dependent cell viability indicates LREX cells in charcoal stripped media (CSS) are resistant to enzalutamide, which is overcome by adding ipatasertib. **c**, Flow cytometry analysis of cell cycle revealed that ipatasertib alone induces cell cycle arrest at the G0/G1 phase and there is an increased sub G1 population when combined with enzalutamide. **d**, Apoptosis analysis using Annexin V/PI dual stain assay indicates increased apoptosis in cells treated with both ipatasertib and enzalutamide.

To determine the mode of cell death following ipatasertib alone or when combined with enzalutamide, we performed cell cycle analysis using propidium iodide (PI) after 24 hours of treatment. Single agent ipatasertib induced a G0/G1 cell cycle arrest (Fig. 2c), and when combined with enzalutamide we found an increased subG1 population, indicative of apoptosis (Fig. 2c). We confirmed the apoptotic cell death using annexin V/PI staining of live cells. Indeed, we observed increased apoptosis in cells treated with both ipatasertib and enzalutamide compared to single agents (Fig. 2d). Taken together, our data suggest that ipatasertib halts cell growth primarily by inducing cell cycle arrest, and when combined with enzalutamide, it induces apoptotic cell death.

### Inhibition of AKT blocks the induction of GR expression and activity through chromatin landscape reorganization

Induction of compensatory glucocorticoid receptor (GR) expression has been reported in a subset of enzalutamide resistant CRPC patients as a mechanism of resistance to AR blockade ^8^. Given the role of GR in enzalutamide resistance in LREX, we sought to determine the effect of AKT inhibition on GR. We examined expression of the GR protein in LREX cells by western blot analysis following AKT and AR inhibition for two days. LREX cells in androgen-deprived media induced GR protein expression, which ipatasertib completely inhibited (Fig. 3a). LREX exposed to enzalutamide induced even higher expression of GR, which was significantly decreased when also exposed to ipatasertib (Fig. 3a). LNCaP have also been shown to induce GR expression, though over longer-term androgen inhibition ^24^. In LNCaP cells, we observed an induction of GR expression after 7 days of exposure to enzalutamide, which is completely blocked by exposure to ipatasertib (**Extended Data Fig. 2a**).

**Fig. 3.**
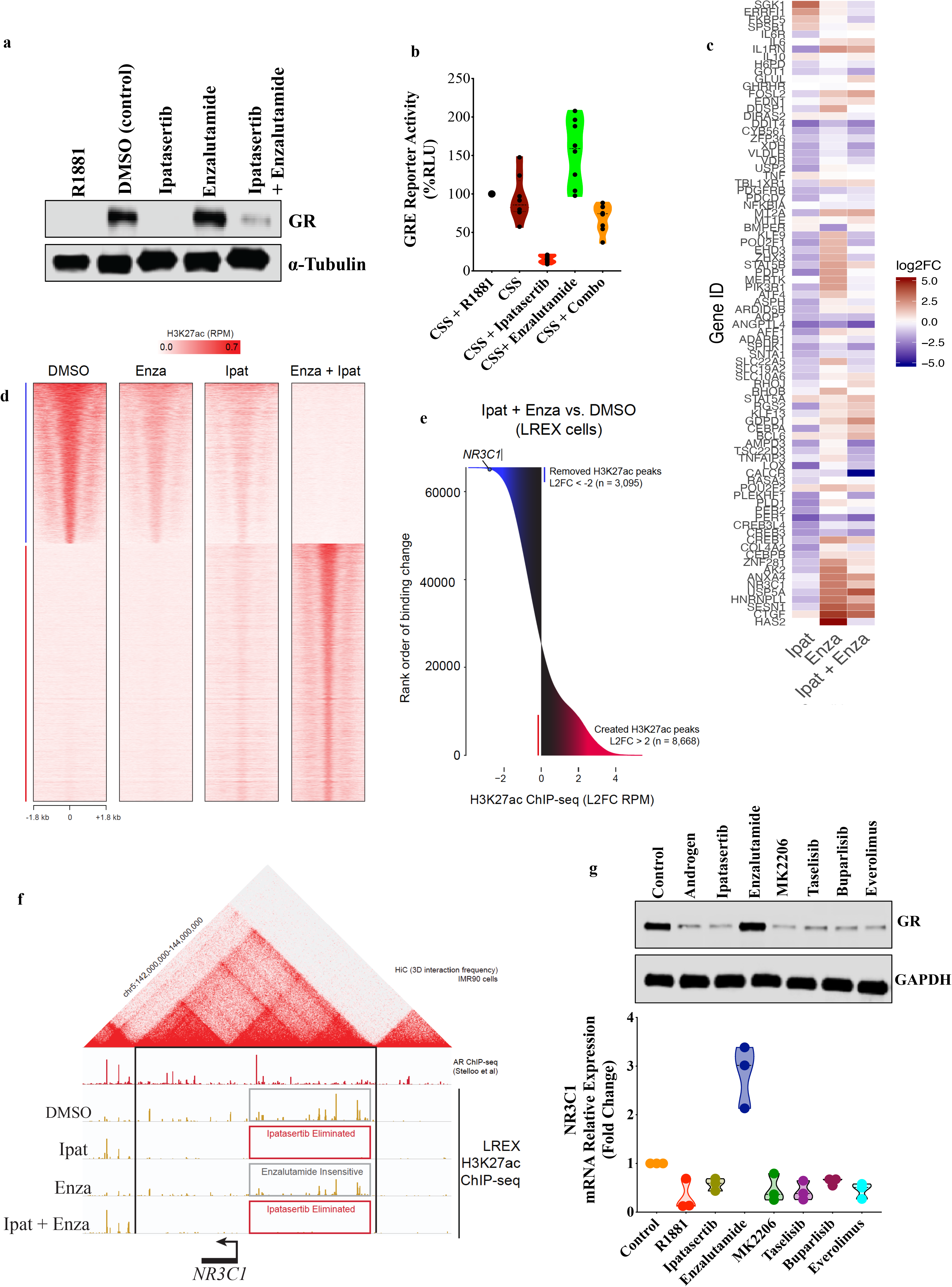
AKT inhibition blocks induction of GR expression and activity. **a**, Western blot indicates GR expression is induced in LREX cells in the setting of 2 days of androgen deprivation or enzalutamide and inhibited by the pan-AKT inhibitor ipatasertib. **b**, Luciferase production from a reporter driven by tandem GRE elements in LREX cells demonstrates decreased GR activity with ipatasertib and increased GR activity with enzalutamide. **c**, Grid heatmap of PCR array evaluating putative GR target genes indicating decreased expression of GR regulated genes with ipatasertib and increased expression after AR inhibition with enzalutamide. **d**, H3K27ac ChIP signal heatmap across treatment groups. **e**, Ranking order of change in H3k27 acetylation upon treatment with ipatasertib and enzalutamide compared to DMSO. **f**, HiC 3-dimensional interaction frequency pyramid and H3K27ac signal tracks (orange) shows regions upstream of the GR gene NR3C1 have absence of enhancer-interaction with ipatasertib treatment or combination treatment. **g**, Western blot (upper panel) and qRT-PCR analysis (lower panel) demonstrate that induction of GR protein or NR3C1 mRNA in the absence of androgen in LREX cells is blocked by synthetic androgen (R1881), AKT inhibitors ipatasertib or MK-2206, PI3K inhibitors taselisib (GDC-0032) or buparlisib (BKM120), or mTOR inhibitor everolimus, but not enzalutamide.

To determine if the decrease in GR expression correlates with GR activity, LREX cells were transfected with a luciferase reporter whose expression is driven by multiple copies of the GR response element. When these cells were exposed to 100 nM ipatasertib for 48 hours, luciferase production decreased significantly. Conversely, when these cells were exposed to enzalutamide, they had increased luciferase production (Fig. 3b). To evaluate GR activity in a more physiologic setting, we performed a PCR array using 84 putative GR-regulated genes. Confirming the luciferase assay finding, ipatasertib lead to decreased expression of a broad range of GR-regulated genes, indicating decreased GR activity (Fig. 3c).

LREX cells have a dynamic reprogramming system with enzalutamide, as these cells are poised to switch on GR for survival when faced with AR blockade. This dynamic on and off switch suggests there is an epigenetic influence on gene expression modulated via typical enhancers (TE) or super-enhancers (SE). Epigenetic processes such as histone modifications at *cis*-regulatory elements can affect gene transcription independent of their orientation or distance via enhancers ^25^. H3K27ac has been established as an important mark of enhancers which distinguishes between active and inactive regions (active referring to a positive influence on the expression of proximal genes) ^26^, hence, regions with deposits of H3K27ac are often associated with enhanced gene activity ^26,27^. We performed Chromatin Immunoprecipitation (ChIP) experiments for H3K27ac in the presence of DMSO, ipatasertib, enzalutamide or the combination of both ipatasertib and enzalutamide, with RNA-seq performed in parallel. First, we looked at global changes of H3K27ac binding across all treatment groups and observed a gradual switch in H3K27ac bound regions in ipatasertib treatment compared to control or enzalutamide and a complete flip in cells treated with ipatasertib and enzalutamide (Fig. 3d). When we ranked in order of binding change, the GR gene, *NR3C1*, ranked amongst the top genes to have decreased H3K27ac deposits (Fig. 3e). To ensure we searched the same cis-regulatory space available to *NR3C1* in 3-dimensions, we used published HiC data ^28^ identifying chromatin interactions inside the nucleus and found clear boundaries (**black lines**, Fig. 3f) of an insulated neighborhood in which active enhancers could directly influence *NR3C1* ^29^. Interestingly, regions on the *NR3C1* locus with enhancer deposition in the presence of enzalutamide or DMSO were completely eliminated with ipatasertib or combination of both ipatasertib and enzalutamide (Fig. 3f). The epigenetic shutdown of *NR3C1* expression was corroborated by RNA-seq and qRT-PCR (**Extended Data Fig. 2b,c**). Taken together, these data suggest AKT inhibition deactivates GR through *cis*-regulatory elements at the *NR3C1* locus that sense the depletion in PI3K-AKT signal.

To determine if inhibition of GR expression is specific to inhibition of AKT by ipatasertib or occurs with antagonism of other PI3K/AKT pathway members, we exposed LREX cells to the AKT inhibitor MK2206, the PI3K inhibitors taselisib or buparlisib, or the mTOR inhibitor everolimus in the setting of androgen deprivation. Stimulation of AR with the synthetic androgen R1881 led to a decrease in GR expression. In contrast, AR inhibition with enzalutamide increased GR expression. All four inhibitors of the PI3K/AKT pathway decreased GR expression similar to that of ipatasertib (Fig. 3g). Taken together, our results suggest PI3K/AKT inhibitors remodel the chromatin landscape to block the induction of GR expression at the transcript level and resensitize cells to enzalutamide.

### Inhibition of GR is required for sensitivity to ipatasertib in the context of GR-dependent tumor growth

To confirm if re-sensitization of cells to enzalutamide by ipatasertib treatment was through inhibition of GR, we genetically knocked down *NR3C1* in LREX cells using shRNA (Fig. 4a,b). We then treated two independent *NR3C1*-KD lines with enzalutamide. Our data demonstrated ∼50% decrease in cell viability compared to the non-silencing (NS) control cells treated with enzalutamide (Fig. 4c), which phenocopies the effect seen when we treat cells with ipatasertib and enzalutamide.

**Fig. 4.**
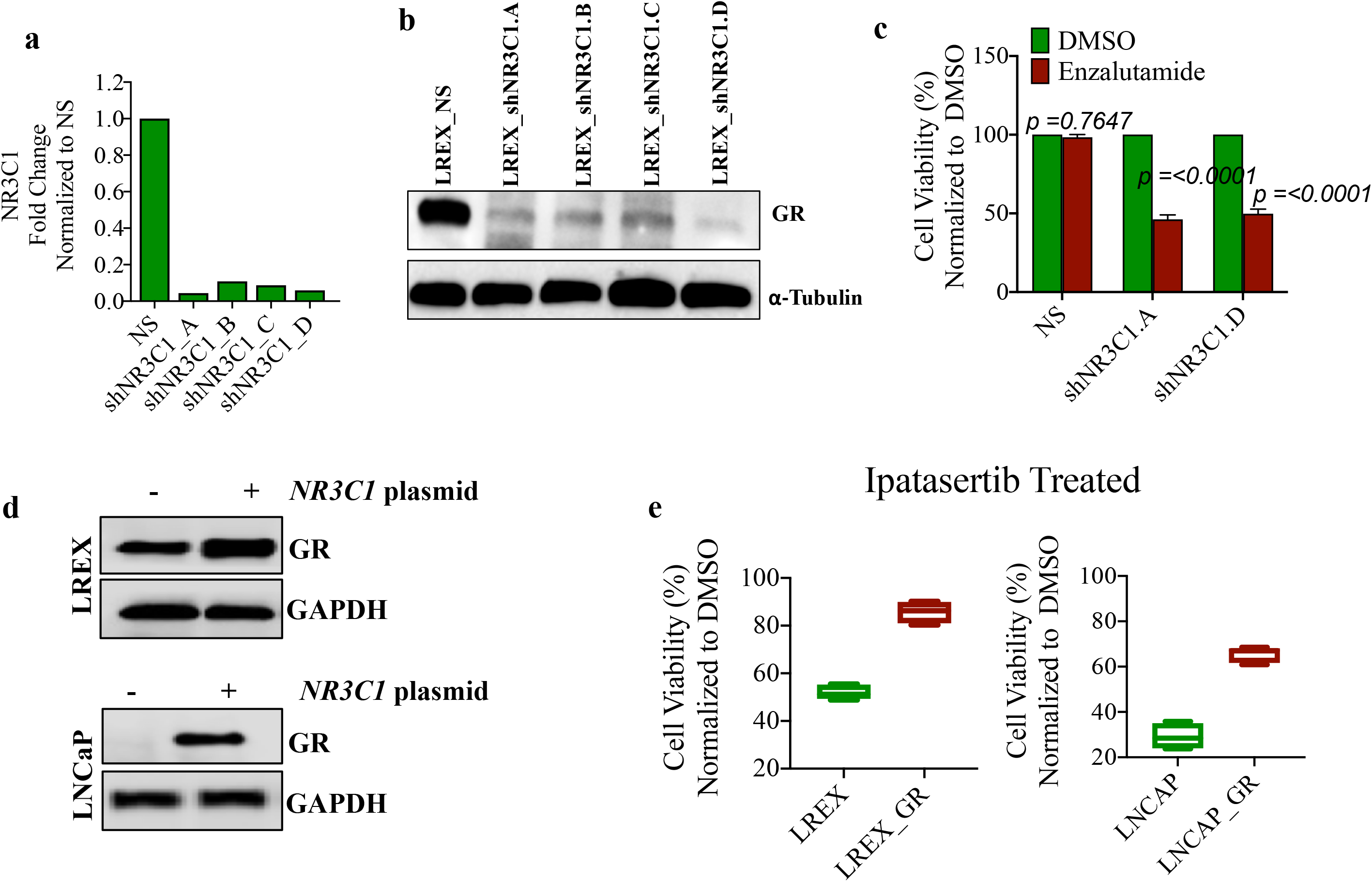
Modulation of GR gene alters responses to enzalutamide and ipatasertib. Efficient knockdown of GR gene (*NR3C1*) and protein in LREX cells as indicated by qRT-PCR, **a** and western blot, **b**. **c**, Cell viability assay indicates sensitization of LREX cells to enzalutamide following *NR3C1* knockdown. **d**, Western blot indicating exogeneous overexpression of GR in engineered LREX (top panel) and LNCAP (bottom panel) cell lines. **e**, Cell viability assay indicates decreased response to ipatasertib in LNCaP and LREX cells overexpressing GR compared to control cells.

Next, we sought to determine if decreased GR expression is required for sensitivity to ipatasertib. We maintained GR expression in LREX and LNCaP cells by transfecting them with a GR-expression construct (Fig. 4d and **Extended Data Fig. 3a,b**). In both models, cells with exogeneous expression of GR demonstrate decreased sensitivity to ipatasertib (Fig 4e). In contrast to LNCaP and LREX, the 22RV1 cell line has high *NR3C1* mRNA and GR protein expression at the basal level and lacks further induction of GR expression following AR inhibition (**Extended Data Fig. 3c, right panel**) ^24^. GR protein level is maintained in the presence of ipatasertib, and the cell line is minimally sensitive to ipatasertib (**Extended Data Fig. 3c, left panel**). This observation suggests that the effect of ipatasertib on GR expression and cell viability is enhanced when GR is induced by AR inhibition, but not when there is constitutive high expression. Overall, our data suggest that AKT inhibition blocks the induction of GR, reducing cell viability and increasing sensitivity to AR blockade.

### Impact of AKT inhibition on GR expression is modulated by AR

We have demonstrated blockade of GR expression at the protein and transcript level using several PI3K/AKT pathway inhibitors, including ipatasertib (Fig. 3g). Interestingly, exogenous androgen (R1881) also suppressed *NR3C1* expression, whereas *NR3C1* expression was increased in the presence of enzalutamide. PI3K-AKT pathway signaling was previously shown to suppress AR activity, and AR is thought to negatively regulate transcription of *NR3C1* ^9^. To determine if AKT inhibition suppresses GR expression and activity through AR-mediated regulation of transcription, we determined the effect of ipatasertib on canonical AR activity. Gene set enrichments analysis (GSEA) was performed using RNA-seq to identify gene sets that were significantly enriched or depleted with each treatment condition. We found among MSigDB Hallmark gene sets, the most upregulated in any condition was Androgen signaling following exposure to ipatasertib (Fig. 5a,b and **Extended Data Fig. 4**). We further examined the expression of a panel of AR-regulated genes following exposure to ipatasertib or enzalutamide. Whereas enzalutamide decreased the expression of AR-regulated genes, ipatasertib caused a mixed response where, on average, the expression of AR target genes was increased (**Extended Data Fig. 5a**). Additionally, we examined AR activity using a luciferase reporter driven by tandem AR response elements. While ipatasertib alone drove similar expression to control, enzalutamide markedly decreased luciferase expression, which was reversed by adding in ipatasertib (**Extended Data Fig. 5b**).

**Fig. 5.**
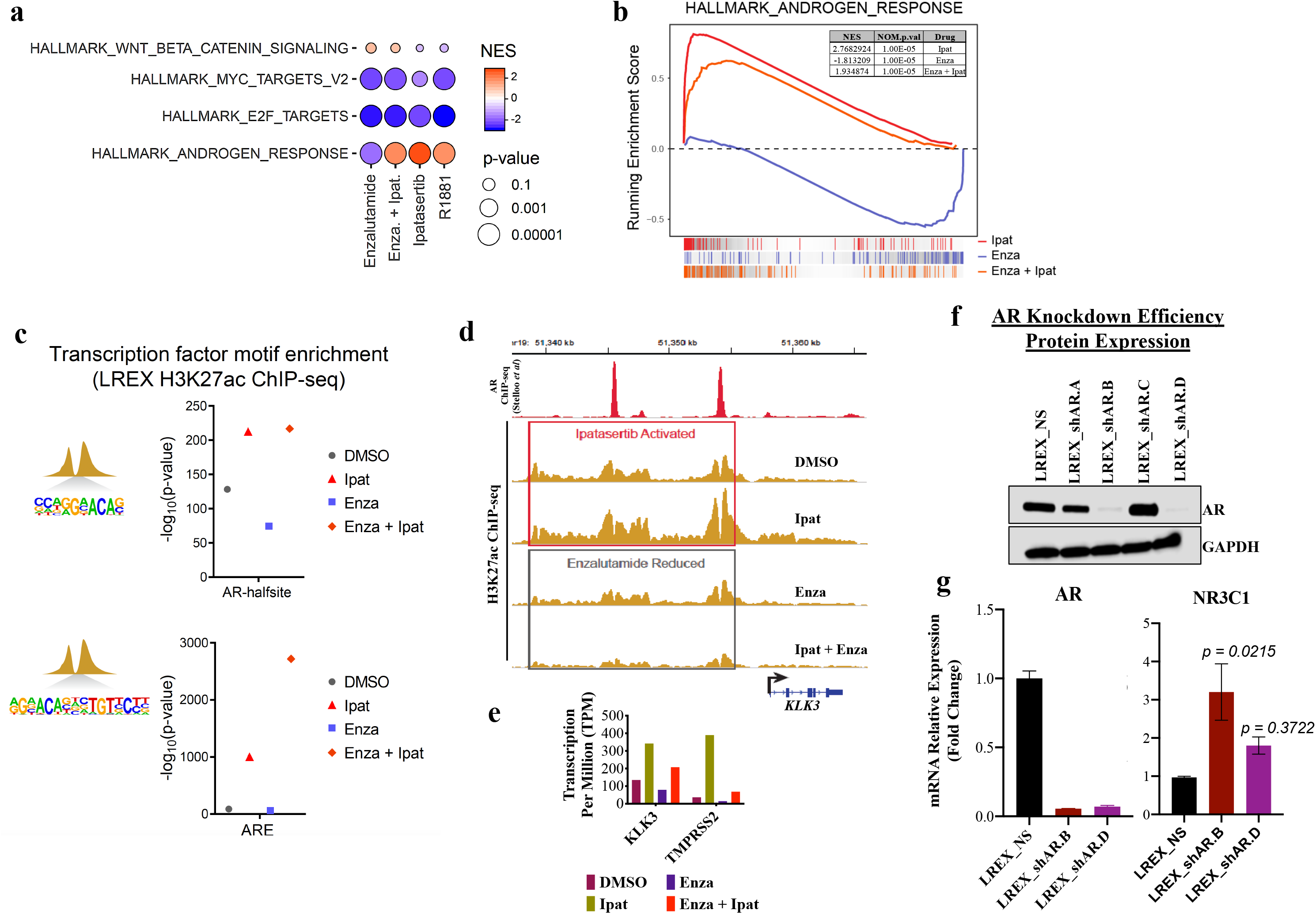
Blockade of GR following ipatasertib exposure is modulated by AR. **a**, Bubble lattice plot of selected enriched hallmark pathways (also see **extended Data fig. 4**). **b**, GSEA comparing gene expression of LREX cells treated with ipatasertib or combination show positive enrichment of genes associated with androgen response and negative enrichment with enzalutamide. **c**, Motif enrichment plots for H3K27ac sites indicates enrichments for AR-half sites and AREs in active promoters and enhancers in LREX cells following exposure to ipatasertib alone or with enzalutamide. **d**, H3K27ac ChIP-seq aligned to published AR ChIP–seq track experiments performed on LREX cells 48hrs post-treatment indicates diminishing of enhancer deposits at the KLK3 locus in cells treated with enzalutamide or combination of both ipatasertib and enzalutamide. **e**, Gene expression data from RNA-seq experiments demonstrates increased expression of the AR regulated genes *KLK3* and *TMPRSS2*. **f**, Western blot indicating successful knockdown of AR in 2 of 4 shAR sequence targets. **g**, QRT-PCR indicates increased *NR3C1* gene expression in AR knockdown cells.

To more broadly examine the effect of ipatasertib exposure on transcription activity, we looked at transcription factor motif enrichment. Motif analysis in H3K27ac-decorated chromatin identified enrichment in samples treated with ipatasertib, both alone and in combination with enzalutamide, of transcription factor sequences recognized by the AR (Fig. 5c). We observed substantial deposition of H3K27ac at several canonical AR target loci, including the *KLK3* super enhancer locus, which was increased by ipatasertib and decreased by enzalutamide (Fig 5d). RNA-seq experiments performed in parallel indicate increased transcription of *KLK3* and *TMPRSS2* with ipatasertib treatment (Fig. 5e).

To further evaluate the direct effect of AR on GR expression at the transcript level, we genetically knocked down the AR gene in LREX cells. We selected two of four target sequences that produced >70% knockdown for our experiments (Fig. 5f). We demonstrate that in both AR-knockdown models, there was an increased in GR expression compared to the non-silencing control, confirming that AR activity directly influences GR expression (Fig. 5g). Taken together, these data indicate AKT inhibition both activates canonical AR targets and blocks GR induction through *cis*-regulatory elements that sense the depletion in PI3K-AKT signal.

### Combination AKT inhibition and androgen inhibition decreases tumor size across multiple xenograft models

To determine how ipatasertib would affect established tumors, either alone or in the setting of AR-targeted therapy, we measured the tumor volume of two independent PDX models treated with ipatasertib and/or castration. These models represent a range of AR activity and AR-responsiveness ^21^. When tumors were well established (∼500 mm^3^), the mice were castrated, given ipatasertib by oral gavage daily (5days on; 2 days off), or both. In both models ipatasertib halted tumor growth as well as or better than castration (average relative change compared to pre-treatment with ipatasertib, −0.3632 vs. control 0.6591; *p=0.002*) (Fig. 6a). Moreover, castration and ipatasertib combined consistently led to decreases in tumor volume in each model (average relative change compared to pre-treatment in combination, −2.737 vs. control, ipatasertib and castration 0.4069; *p=0.0004*) (Fig. 6a).

**Fig. 6.**
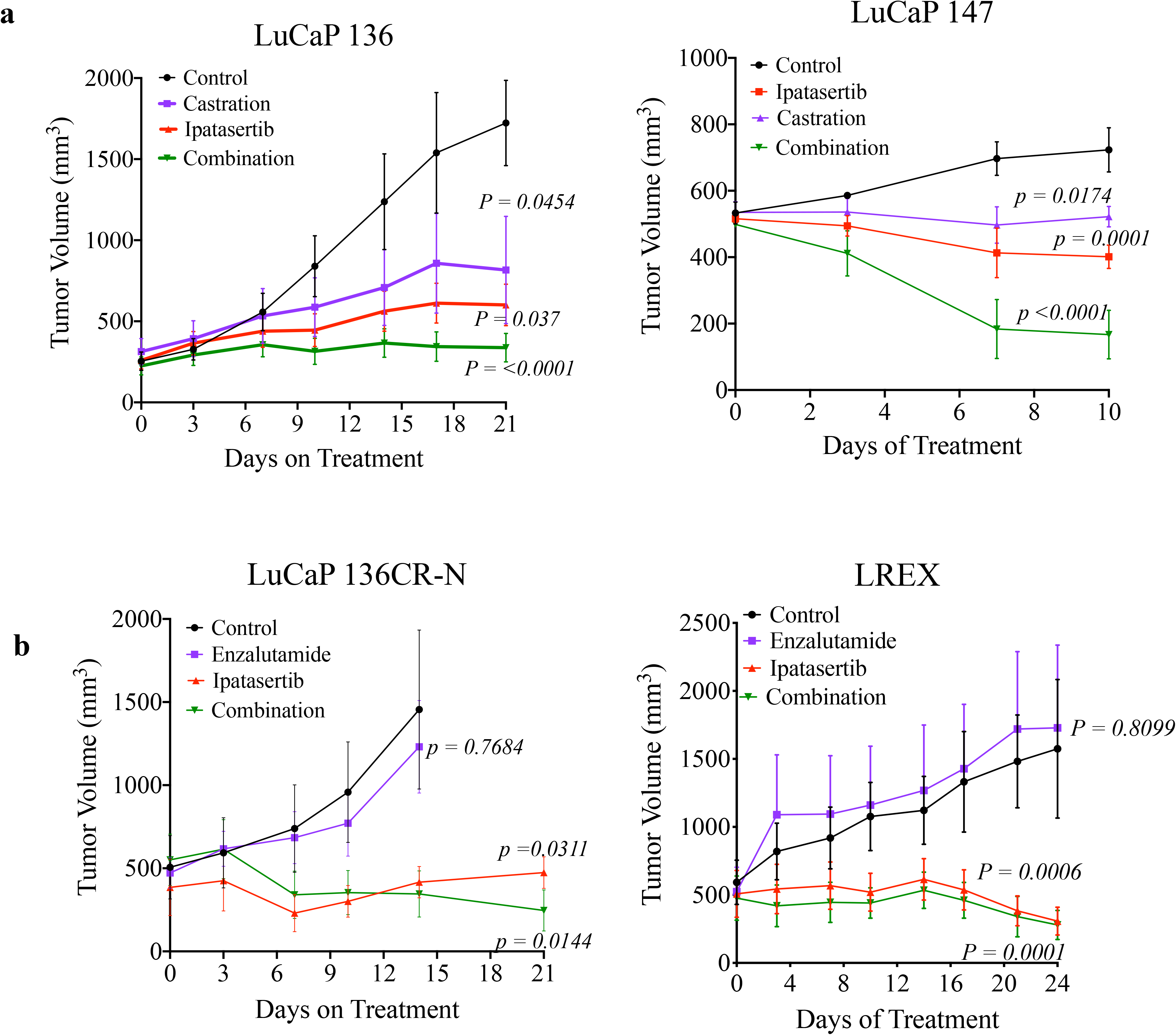
Ipatasertib and AR inhibition combine to inhibit tumor growth in *in vivo* prostate cancer models. **a**, In models of androgen sensitive prostate cancer LuCaP 147 and LuCaP 136 grown in intact mice, ipatasertib decreased tumor burden as monotherapy, with enhanced efficacy when combined with castration. **b**, In enzalutamide resistant prostate cancer LuCaP 136CR-N PDX and LREX grown in castrated mice, ipatasertib effectively inhibits tumor growth as a single agent and in combination with enzalutamide. Graphs are presented as the mean +/− SD. (n=7-8/group for LuCaP 136, 136CR and LREX, n=6/group for LuCaP 147).

We additionally tested ipatasertib and AR inhibition in two models of CRPC. The LuCaP 136CR-N model was derived *in vivo* from the parental LuCaP 136 model grown in castrated mice. We also examined xenografts grown from the LREX cell line. Castrated mice harboring tumors from these CRPC models were exposed to ipatasertib, enzalutamide, or both. In both CRPC models, enzalutamide had little effect, with tumor growth rates similar to that of controls (average relative change compared to pre-treatment in enzalutamide, 0.5180 vs. control, 0.5844; *p=0.99*) (Fig. 6b). Ipatasertib, however, caused a dramatic slowing of tumor growth or a decrease in tumor volume (average relative change compared to pre-treatment with ipatasertib, −14.95 vs control and enzalutamide, *p=0.0003).* The combination of enzalutamide and ipatasertib showed similar results (average relative change compared to pre-treatment with combination, −23.15 vs control and enzalutamide *p=0.0001*) (Fig. 6b). Thus, there is a consistent anti-tumor effect of AKT inhibition in diverse *in vivo* prostate cancer models, representing both hormone sensitive and resistant disease.

### AKT inhibition enhances AR activity and decreases GR expression *in vivo*

Having demonstrated in cell line models that PI3K/AKT pathway inhibitors block GR expression, and that this regulation of GR expression is in part through increased AR activity, we sought to confirm these findings in mouse xenograft models. We took end of treatment tumors from experiments shown in Figure 6 and evaluated mRNA and/or protein of GR and AR-regulated genes. Similar to in vitro experiments, castration or enzalutamide led to a marked decrease of AR activity, as indicated by decreased expression of downstream genes *KLK3* and *NKX3-1*, whereas ipatasertib caused increased expression of these genes (Fig. 7a,b,c,d). Ipatasertib caused concomitant decrease in GR protein level, which was maintained or increased with castration (Fig. 7e). Similar results were seen in CRPC models treated with enzalutamide (Fig. 7f). These effects were maintained throughout treatment, as tumors were harvested after 2 or 4 weeks of therapy.

**Fig. 7.**
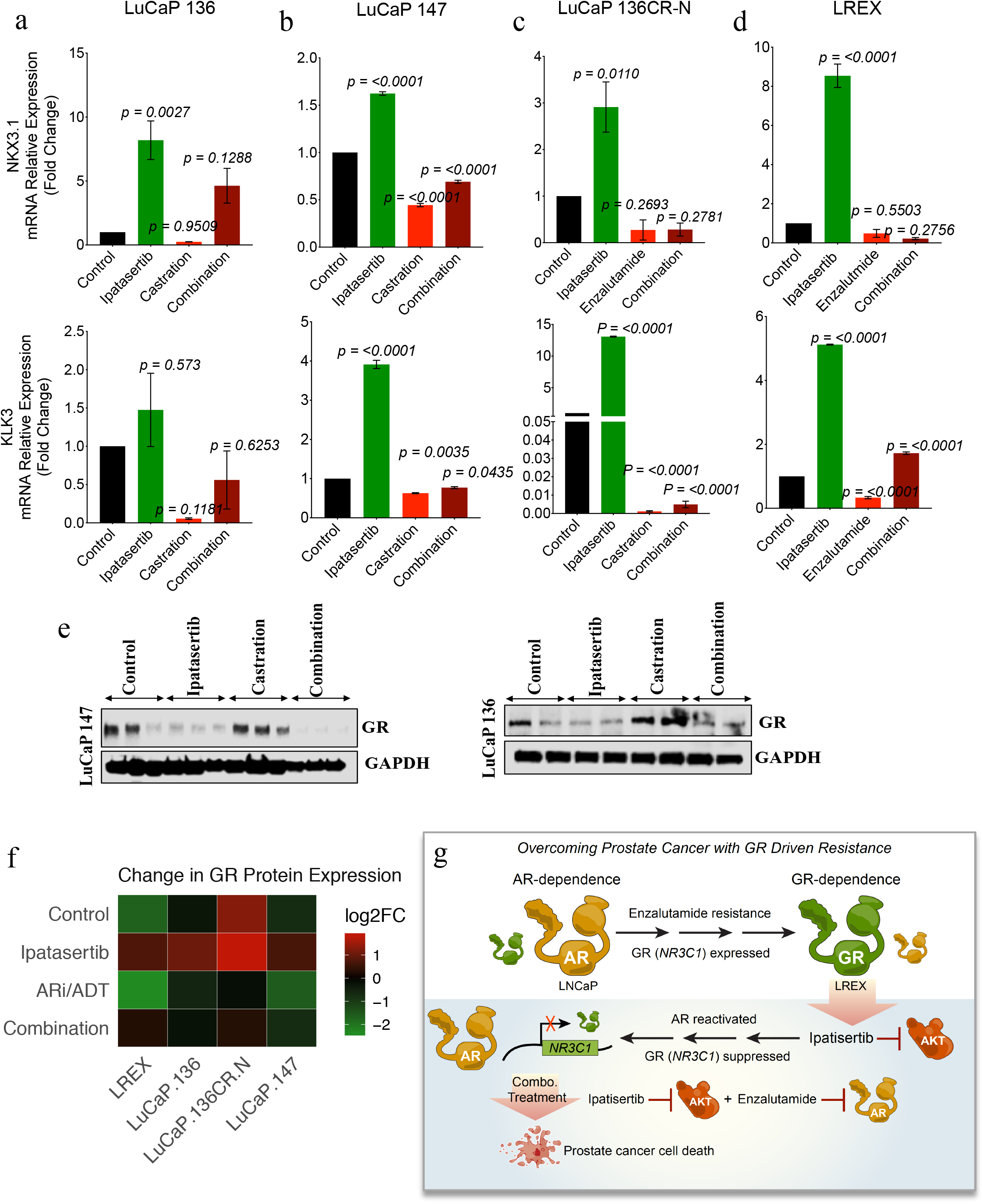
AKT inhibition enhances AR activity and blocks GR expression in vivo. **a-d**, RT-PCR from 3 patient-derived xenografts and one cell line treated with the pan-AKT inhibitor ipatasertib (100 mg by oral gavage), castration, enzalutamide or combination of ipatasertib + castration/ ipatasertib + enzalutamide. As expected, expression of canonical AR-regulated genes *KLK3* (PSA) and *NKX3-1* are decreased with castration or enzalutamide. Expression of these genes is increased with ipatasertib or with enzalutamide combined with ipatasertib. **e**, Representative western blot analysis from LuCaP 136 and LuCaP 147 showing increased GR expression following castration, and decreased levels after exposure to ipatasertib. **f**, Summary heatmap of all models shows the expression of GR protein is decreased with ipatasertib, which is attenuated modestly in castrated/enzalutamide treated animals in most models. **g**, Model of regulation of GR by AKT inhibition in the context of GR-dependent growth. AKT inhibition decreases GR at the transcriptional level through induction of AR activity. Graphs are presented as mean +/− SEM.

## DISCUSSION

The PI3K/AKT pathway is a complex signaling pathway that is involved in survival, proliferation, metabolism, and growth pathways. In the current study we show that inhibition of the PI3K/AKT signaling pathway can block GR expression that is induced as a resistance mechanism to AR-targeted therapy, both when combined with androgen deprivation or the AR antagonist enzalutamide. In multiple prostate cancer xenograft models representing a spectrum of AR-dependence, this leads to significant anti-tumor response. Blockade of GR expression is associated with reorganization of the chromatin landscape, induction of canonical AR activity, and decreased expression of the GR gene, *NR3C1* (Fig. 7g).

While multiple mechanisms can lead to resistance to AR-targeted therapy, one that has drawn particular interest is the induced expression of GR, leading to a new therapeutic strategy for a subset of prostate cancer patients. Interestingly, the upregulation of GR as compensatory hormone receptor signaling has also been reported in breast cancer and is associated with poor prognosis in triple-negative breast cancer ^30,31^. It is likely that diverse mechanisms can regulate GR expression in diverse prostate cancers subtypes. Indeed, 22Rv1 cells, which express constitutively high levels of GR that is not affected by AR inhibition nor ipatasertib, are relatively resistant to ipatasertib. Nevertheless, we demonstrate here that ipatasertib blocks the induction of GR expression across numerous models tested, both *in vitro* and *in vivo*, which is associated with ipatasertib sensitivity. In two engineered GR-overexpressing cells lines we show enforced GR expression reduces ipatasertib sensitivity, demonstrating the key role of GR expression and activity in response and resistance to AKT inhibition. To our knowledge, this is the first demonstration that PI3K/AKT pathway inhibition blocks GR expression in prostate cancer, leading to significant anti-tumor effect.

The strongest evidence to date for upregulation of GR is in response to enzalutamide for mCRPC ^8,9,24,32^. In advanced prostate cancer ipatasertib has been used in combination with abiraterone, which is given with prednisone, providing further rationale for blockade of GR contributing to anti-tumor effect. The importance of GR expression for development of castrate resistance in newly diagnosed metastatic disease in response to initial therapy, whether with ADT alone, ADT and docetaxel, or ADT and abiraterone/prednisone, has not been investigated. It is worth noting that we found marked anti-tumor effect with ipatasertib monotherapy and in combination with castration in multiple PDX models of HSPC. In some respects, high risk localized disease may have similar biology to metastatic HSPC. Upregulation of GR expression has been demonstrated in a subset of patients in both neoadjuvant enzalutamide and neoadjuvant abiraterone/prednisone trials in this setting ^33,34^, raising the question of whether adding a PI3K/AKT inhibitor could improve the outcomes for those with HSPC. One explanation for the prevalence of GR expression in these early phase studies, as opposed to well-characterized genomic alterations (e.g. AR enhancer amplification), is that development of resistance through selection of genomically altered subclones may require more time, and later stage disease, than induction of GR.

In our studies regulation of GR induction by the PI3K/AKT pathway is associated with remodeling of the chromatin landscape. It is likely there are additional important consequences of chromatin remodeling aside from regulation of GR expression and activity. The consistent effects on GR across diverse models, however, and the requirement for GR inhibition for maximal inhibitor sensitivity, suggest that the effect on the *NR3C1* gene and GR activity may be crucial. The PI3K/AKT pathway has been demonstrated to have effects on the chromatin landscape in breast cancer, as well, another hormone receptor-regulated cancer ^35^. The full consequences of AKT inhibition on chromatin remodeling in prostate cancer have yet to be elucidated.

Along with inhibition of GR, AKT inhibition induces an increase in canonical AR activity. This seems counterintuitive, since many prostate cancer therapies are designed to inhibit AR activity. Yet others have noted in tumor samples that high AR activity is associated with low cell proliferation ^36^. It is not clear if this is the same mechanism behind responses induced by supraphysiologic testosterone in recent clinical trials ^37,38^. It may be that increased canonical activity is associated with decreased non-canonical activity. More extensive studies will be required to tease out the paradoxical mechanisms behind different levels of AR activity.

It has been suggested that PI3K/AKT targeted therapies might drive differentiation toward t-SCNC, as demonstrated with the LNCaP cell line ^39^. In our data, however, a PDX model of small cell cancer (LuCaP 145.1) was particularly sensitive to AKT inhibition in 3D organoid culture (**Extended Data Fig. 6**). Another responsive model *in vivo*, LuCaP 136, represents aggressive variant prostate cancer due to loss of function of both PTEN and TP53 ^40^. Indeed, others have demonstrated that activated AKT and N-MYC combine to drive a neuroendocrine phenotype ^41^, supporting the use of an AKT inhibitor for tumors with neuroendocrine features.

This study was focused on prostate cancer, for which induction of GR expression is a demonstrated mechanism of resistance for established therapies. PI3K-pathway inhibition in general, and AKT inhibition with ipatasertib in particular, have been and continue to be tested in a number of other solid tumors. In breast cancer, GR expression has been shown to play a role in resistance to taxanes, with apparently opposite effects in hormone receptor positive and hormone receptor negative disease ^30^. It is not known if ipatasertib blocks GR activity in breast cancer as it does in prostate cancer, or if that plays a role in the efficacy seen in a recent phase II study ^16^.

The long history of PI3K/AKT pathway inhibitors for the treatment of prostate cancer and other solid tumors has translated to only modest success in the clinic. The pan-AKT inhibitor ipatasertib shows promise in combination with abiraterone plus prednisone for late stage prostate cancer. Our data demonstrate marked anti-tumor effect in models that upregulate GR to induce resistance. A better understanding of the clinical settings in which GR activity is most critical will help usher in a new target in the prostate cancer therapy armamentarium.

## Conflict of interest

The authors declare no conflict of interest.

## ACKNOWLEDGMENT

We would like to thank Dr. Charles Sawyers, Memorial Sloan Kettering Cancer Center, for providing LREX cells, and Dr. E. Corey and Dr. R. Vessella, University of Washington, for providing the LuCaP models. We are grateful to the National Cancer Institute (NCI) Developmental Therapeutics Program for providing ipatasertib and enzalutamide used for animal studies, and the Illumina Sequencing Facility, Center for Cancer Research, for assistance with library preps and next generation sequencing. This research was supported by the Intramural Research Program, NCI, NIH, the Prostate Cancer Research Program under Award No. W81XWH-13-1-0451 (DVW), and the Prostate Cancer Foundation (DVW).

