## Supplementary Table1, Supplementary Fig1-6 for "Targeting the PI3K/AKT pathway overcomes enzalutamide resistance by inhibiting induction of the glucocorticoid receptor"

### Supplementary Figures

### Extended Figure Legends

**Supplementary Table 1**. Summary of primer sequences and constructs.

**Supplementary Figure 1. Assessment of ipatasertib as single agent across models representing hormone sensitive PCa in vitro.** (A) Graph presentation of cell viability at 48hrs in androgen sensitive PTEN null (LNCAP) and PTEN intact (LAPC) cells indicates significant decreases in viable cells treated with 100 nM pan-AKT inhibitor ipatasertib. (B) Western blots in tested cell lines indicates attenuation of AKT signaling by the ATP-competitive inhibitor ipatasertib, correlating with increase in AKT phosphorylation and decreased phosphorylation of downstream targets.

**Supplementary Figure 2. Assessment of ipatasertib on GR expression.** Supplementary Figure 2. Assessment of ipatasertib on GR expression. (A) Western blot analysis (right) and qRT-PCR mRNA analysis indicates LNCaP cells induce GR expression in the presence of enzalutamide for 7 days, which is blocked by the pan-AKT inhibitor ipatasertib. (B) RNA-Seq data demonstrates decrease GR expression at the transcriptional level with ipatasertib treatment. (C) qRT-PCR data validating NR3C1 gene expression. ****p>0.0001, ns=not significant

**Supplementary Figure 3.** Quantitative RT-PCR and western blot indicating exogeneous overexpression of GR in engineered LNCAP (A) and LREX (B) cell lines. (C) 22RV1 cells have constitutively high GR expression that is not decreased with ipatasertib (right panel) and have marginal response to ipatasertib (left panel).

**Supplementary Figure 4. B**ubble-lattice plot indicating MSigDB Hallmark gene sets

**Supplementary Figure 5.** Inhibition of AKT enhances AR activity (A) Heatmap of PCR array evaluating AR target genes indicates heterogeneity in expression of AR target genes; violin plot summarizing expression of genes indicating overall increased expression with AR regulated gene by AKT inhibition, and decrease with enzalutamide compared to control (CSS + DMSO). (B) Luciferase production from reporter driven by tandem ARE elements in LREX cells demonstrate decreases expression with AKT inhibition and increased expression with enzalutamide. Results are presented in mean + SEM

**Supplementary Figure 6.** High-through-put 3 dimensional organoid drug screens for Ipatasertib indicates heterogenous response to across patient derived xenograft organoids.

### Extended Table 1.

| Item ID | Sequence |
|---|---|
| AR | |
| Forward Primer | GGTGAGCAGAGTGCCCTATC |
| Reverse Primer | GCAGTCTCCAAACGCATGTC |
| KLK3 | |
| Forward Primer | CACACCCGCTCTACGATATG |
| Reverse Primer | GAGGTCCACACACTGAAGTT |
| NKX3-1 | |
| Forward Primer | AGCCAGAAAGGCACTTGG |
| Reverse Primer | TCACCTGAGTGTGGGAGAA |
| NR3C1 | |
| Forward Primer | TCTGAACTTCCCTGGTCGAA |
| Reverse Primer | GTGGTCCTGTTGTTGCTGTT |
| GAPDH | |
| Forward Primer | AGT CCC CAG AAA CAG GAG GT |
| Reverse Primer | AGAGCGCGAAAGGAAAGAA |
| MYCOPLASMA | |
| Forward Primer | TGCCTGGGTAGTACATTCGC |
| Reverse Primer | GCGGTGTGTACAAACCCCGA |
| ShAR (OriGene) | |
| A | GGCTCCAGCGACAGCCAACCGCCTCTTGCA |
| B | GTCCTGGATGAGGAACAGCAACCTTCACA |
| C | ACAACGCCAAGGAGTTGTGTAAGGCAGTG |
| D | GCTGACAGTGTCACACATTGAAGGCTATG |
| shNR3C1(OriGene) | |
| A | ACTGGCTGTCGCTTCCTCAATCAGACTCCA |
| B | GCGAGCGGCTCCTCTGCCAGAGTTGATAT |
| C | CTGGTGTCACTGTTGGAGGTTATTGAACC |
| D | CCAGTGAGATTACAGAGGAAGTTATCCTC |

**Extended Figure 1. Assessment of ipatasertib as single agent across models representing hormone sensitive PCa in vitro.**

A

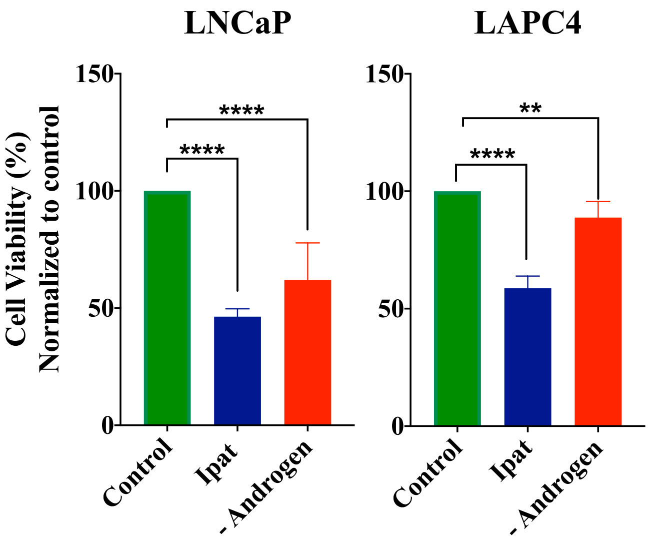


B

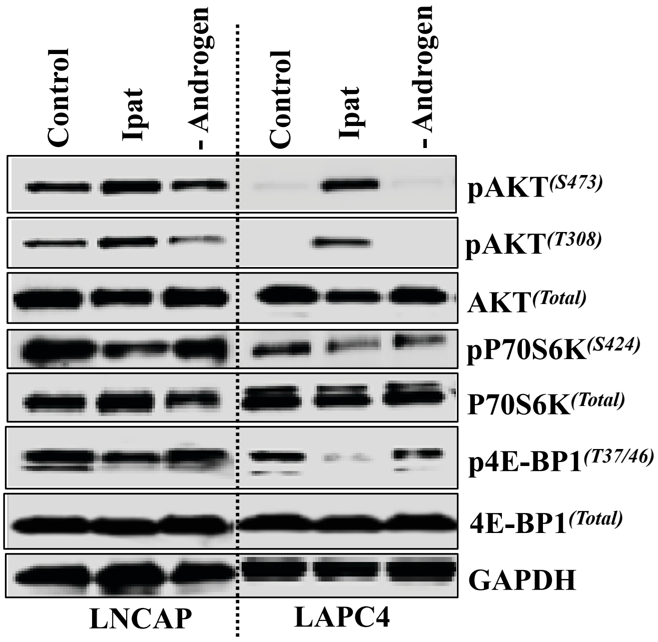

**Extended Figure 2. Assessment of ipatasertib on GR expression.**

a

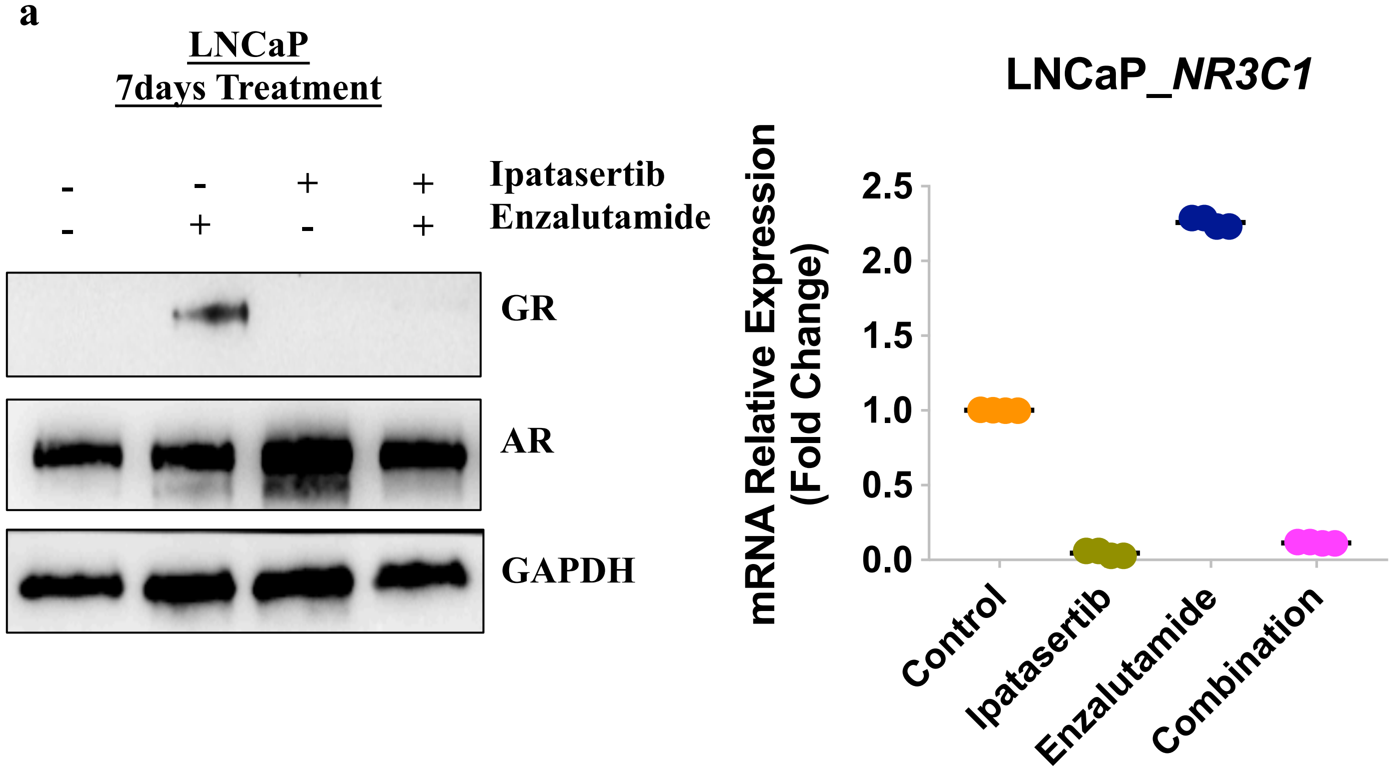


b

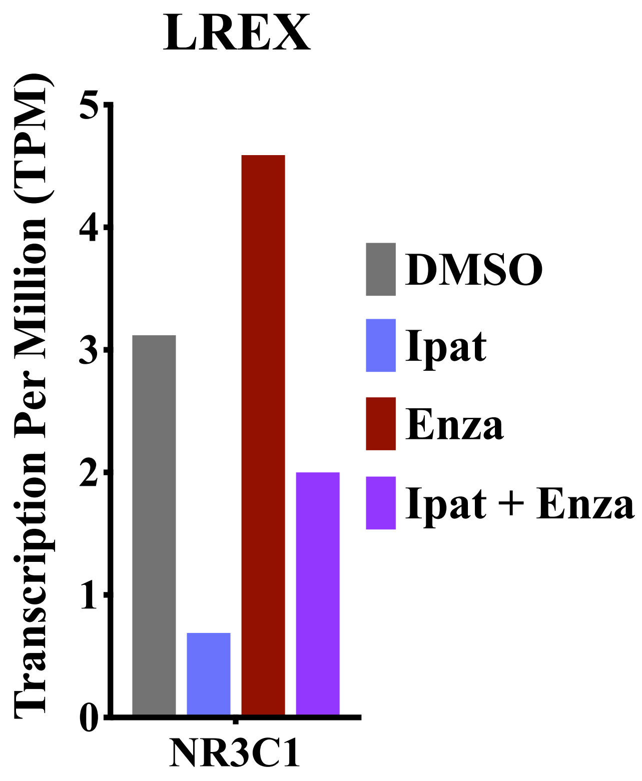


c

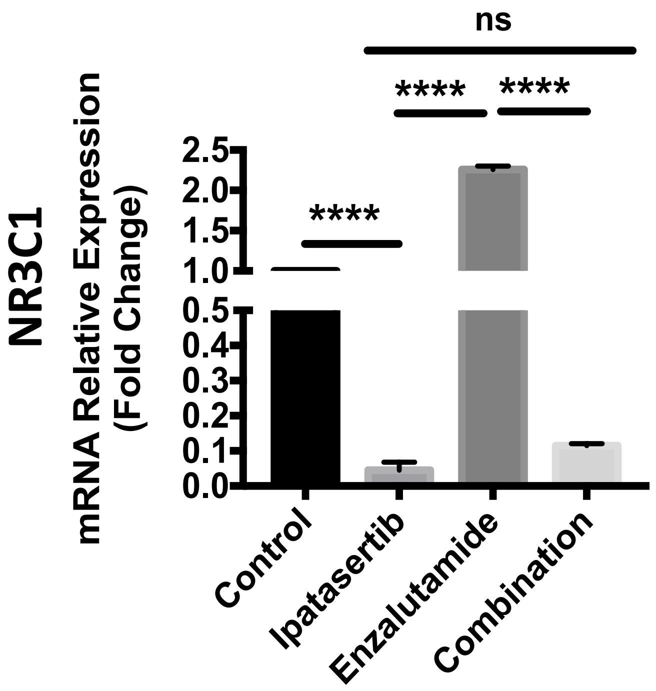

**Extended Figure 3: Quantitative RT-PCR and western blot indicating exogeneous overexpression of GR in engineered LNCAP and LREX cell lines**

a

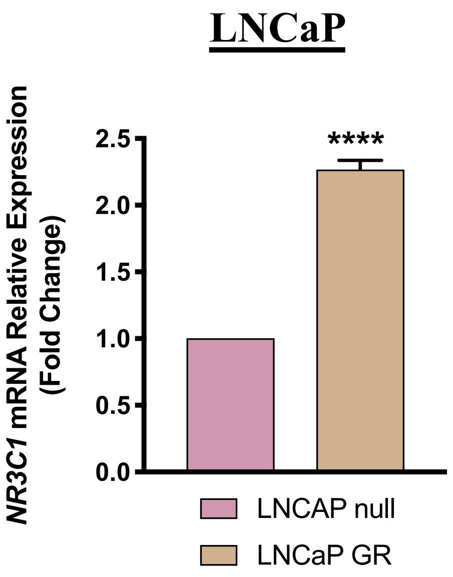


b

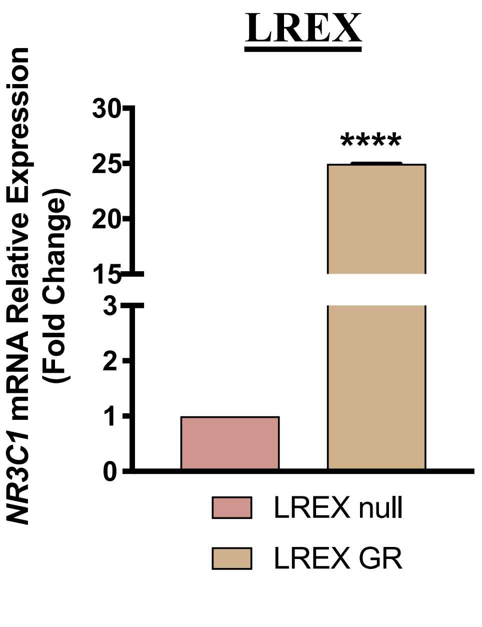


c

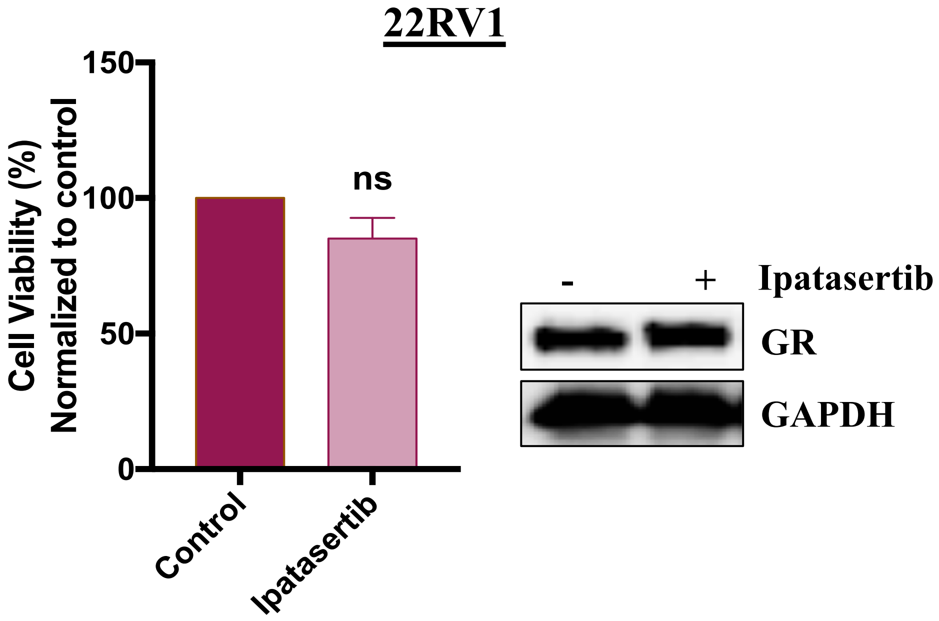

### Extended Figure 4

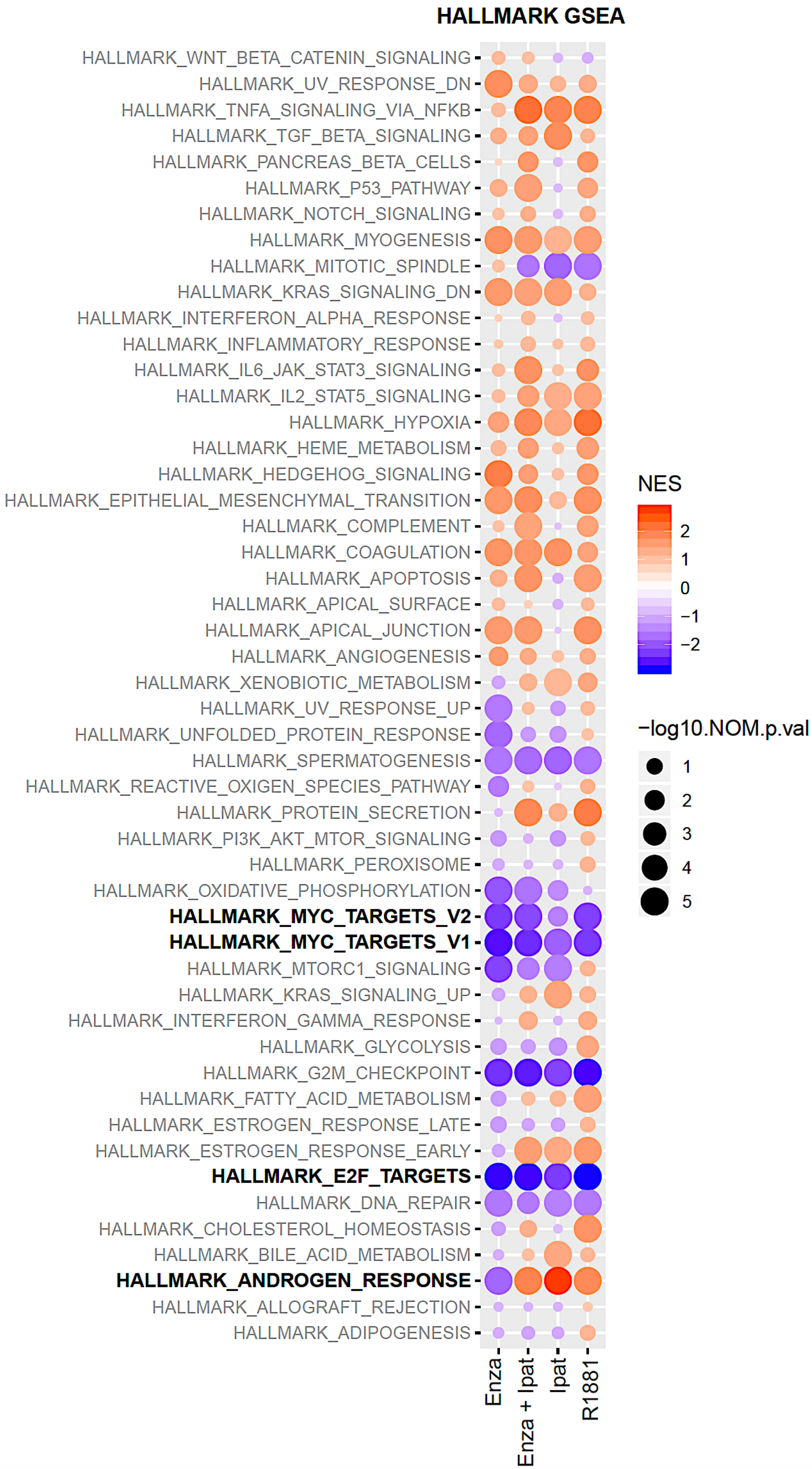

### Extended Figure 5: Inhibition of AKT enhances AR activity

**a**

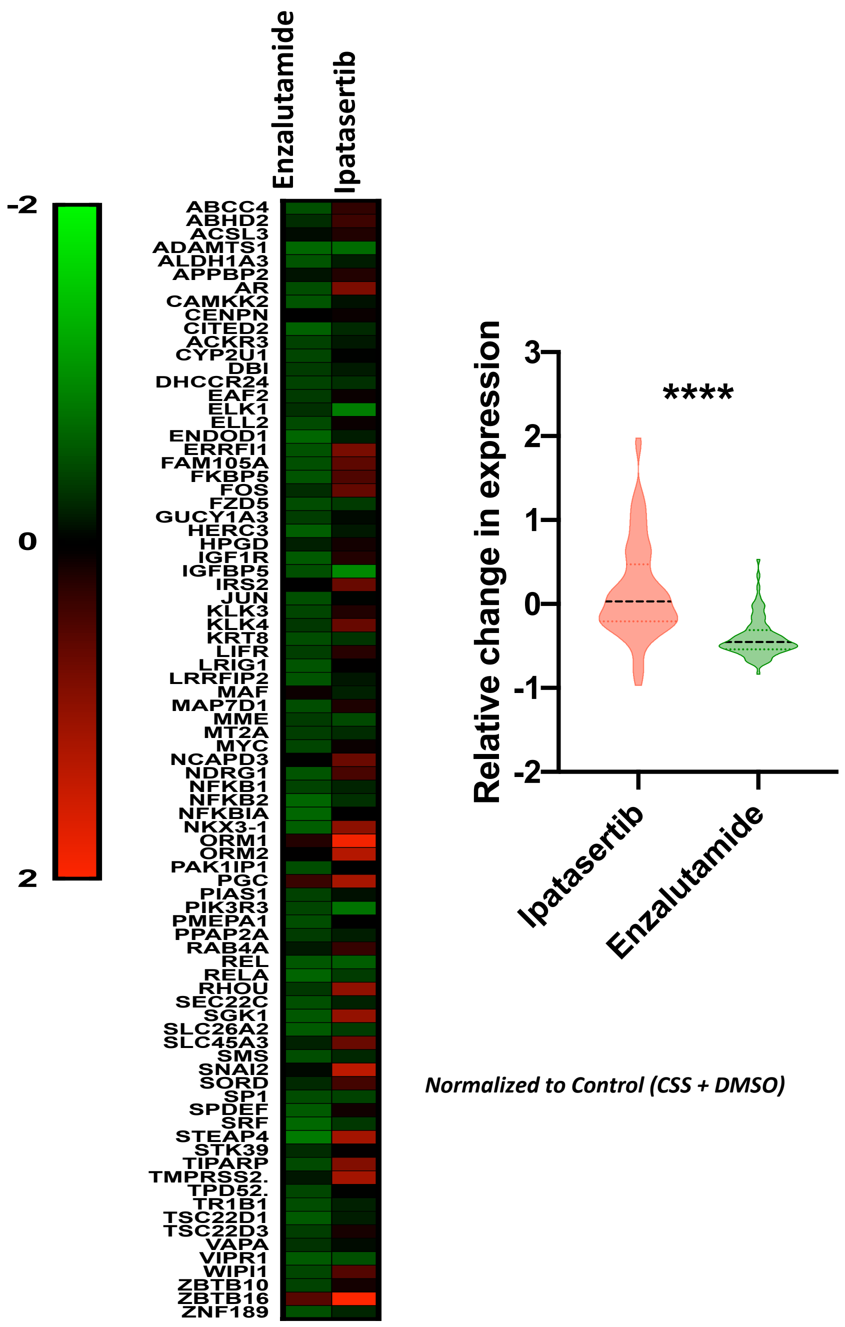


**b**

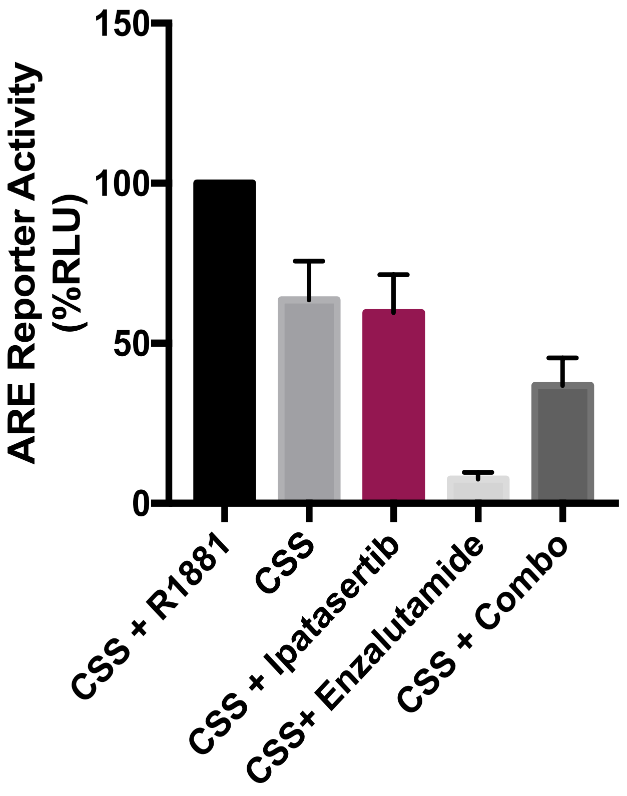

**Extended Figure 6: High-through-put 3 dimensional organoid drug screens for Ipatasertib indicates heterogenous response to across patient derived xenograft organoids**

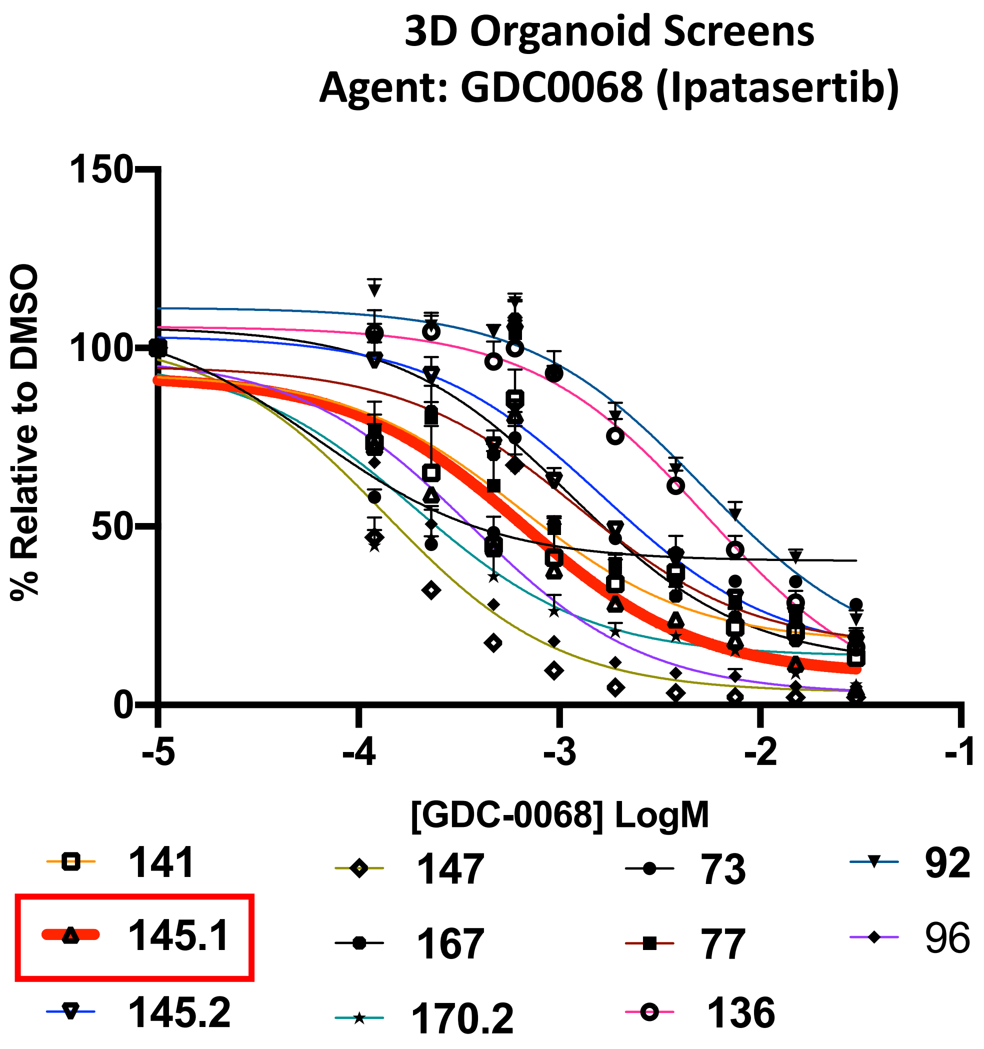
